# Prediction of Systemic Lupus Erythematosus from Genotype and Gene-Expression Data: Batch Correction and Leakage-Free Evaluation

**DOI:** 10.64898/2026.10.18.759998

**Authors:** Vasudha Kannan, R Karthik, Aldis Daniel, Hitesh Kandarpa

## Abstract

Systemic Lupus Erythematosus (SLE) is a complex autoimmune disease with a strong genetic component and highly variable clinical features. Because of this heterogeneity, predicting disease status remains challenging using any single marker alone. In this study we explore two complementary data sources, (1) genotypes at 30 candidate SNPs in a South Indian case–control cohort (*n* = 425) and (2) bulk expression of 13 genes from the same pathways, curated from six GEO series (*n* = 642 SLE or control samples from 483 individuals), to investigate whether they can be used to distinguish SLE cases from healthy controls. The expression data were placed on a common log_2_ scale and corrected for batch effects with mean-only ComBat. To avoid data leakage, ComBat, gene filtering, scaling and model fitting were re-estimated inside every training fold, and all samples from the same individual were kept in the same fold. Performance was summarised with ROC-AUC, precision–recall AUC (PR-AUC) and the Matthews correlation coefficient (MCC) over 10 *×* 5-fold cross-validation, the models were interpreted with out-of-fold SHAP values, and the whole pipeline was checked with label permutations. For the SNP data, a dominant encoding with age and sex gave ROC-AUC 0.699 *±* 0.052, PR-AUC 0.668 *±* 0.063 and MCC 0.309 *±* 0.101, but this depended on four SNPs that deviate from Hardy–Weinberg equilibrium in controls; without them the SNPs added little to age and sex (ROC-AUC 0.656). The strongest single association, at *MTRR* A66G, came from an excess of heterozygotes among controls, a pattern typical of genotyping error, whereas homozygous *CYP1A1* M2 and *COMT* genotypes and *CYP1A1* M4 carriers were enriched in cases, pointing to xenobiotic metabolism. The leakage-free expression classifiers reached ROC-AUC 0.837 *±* 0.033 (L1-penalised logistic regression) and 0.915 *±* 0.021 (XGBoost), and shuffled labels gave chance-level results, confirming that the pipeline does not leak. However, a model given only the study label reached ROC-AUC 0.857, because disease status is strongly confounded with study. Within individual studies the ROC-AUC was 0.743 (L1-LR) and 0.835 (XGBoost), and it fell to 0.46–0.74 when whole studies were held out. Lower *MTR* expression was the most important and most consistent expression feature, whereas two other highly ranked genes, *CYP1A1* and *GSTT1*, carried little signal within studies. For pooled multi-study data, leakage-free cross-validation alone does not guarantee a reliable estimate; batch–label confounding and genotype quality have to be checked directly.

## 1 Introduction

Systemic Lupus Erythematosus (SLE) is a systemic autoimmune disease characterised by loss of tolerance to self-antigens, auto-antibody production and immune-mediated tissue damage that can affect multiple organs, including the skin, joints, kidneys and central nervous system (Tsokos, 2011). The accurate classification of SLE is clinically important because it influences the intensity of treatment, organ-specific monitoring and prognosis, yet current criteria are still largely based on clinical and serological features rather than molecular data (Aringer et al., 2019). This motivates the development of data-driven machine-learning approaches that can use genetic and gene-expression information to refine disease-state prediction and improve patient stratification (Figgett et al., 2019).

In this work, we adopt a dual approach to predict disease state using both genotype data and bulk gene-expression data. By analysing these two modalities together, we aim to capture both inherited risk signals and the downstream transcriptional changes associated with SLE.

Public expression data make this kind of study possible, but pooling them brings two problems. First, samples collected in different studies and on different microarray platforms carry strong batch effects, and per-gene standardisation does not remove them (Chen et al., 2011). Second, any preprocessing step that is fitted on all samples before cross-validation, or repeated measurements of the same person that end up in both the training and the test fold, can leak information from the test set into training and inflate performance (Kapoor and Narayanan, 2023). An earlier version of this analysis normalised the expression data with per-gene z-scores across the pooled cohort and reported ROC-AUC, F1 and accuracy on folds split by sample. In this version we (i) correct batch effects explicitly with ComBat (Johnson et al., 2007), (ii) rebuild the evaluation so that no step sees the test fold and no individual appears on both sides of a split, and (iii) report precision–recall AUC and the Matthews correlation coefficient, which are more informative than ROC-AUC and accuracy when the classes are imbalanced (Saito and Rehmsmeier, 2015; Chicco and Jurman, 2020). We also measure how much of the expression signal can be explained by study membership alone, explain both models with out-of-fold SHAP values and per-SNP effect sizes, and test the whole evaluation pipeline with label permutations.

We tested four hypotheses:

- **H1:** Dominant encoding outperforms additive encoding for SNP-based classification.
- **H2:** Folate-pathway SNPs show the strongest association with SLE.
- **H3:** XGBoost will not substantially outperform logistic regression given the small sample size.
- **H4:** Gene-expression data will be more discriminative than SNP data.

## 2 Datasets

### 2.1 Genotype data

The SNP genotype data set was taken from Rupasree et al. (2016), a study designed to assess the contribution of genetic variation at the pathway level to the susceptibility to SLE in a South Indian cohort. It comprises 188 SLE-diagnosed individuals (178 females, 10 males) and 237 healthy controls (230 females, 7 males), totalling 425 subjects. Genotypes at 30 SNP loci were measured using PCR-based methods and are encoded as 0, 1 or 2, representing the number of copies of the alternate allele present (homozygous reference, heterozygous and homozygous alternate respectively). The SNPs were selected from the folate, xenobiotic-metabolism, Toll-like receptor and STAT4 pathways studied by Rupasree et al. (2016). In addition, the data include age at sample collection and biological sex. No subject appears more than once. The genotype codes follow the source data. For several SNPs the genotype coded 2 is the most common one in this cohort (for example *PTPN22* rs2476601, *IRF5* rs2280714 and *STAT4* rs7574865), and four markers (*MTHFR*, *GST T1*, *GST M1* and the *TNF α* CCC haplotype) take only the values 0 and 1. The allele behind each code was not supplied with the data, so effect directions in this paper are reported relative to this coding.

### 2.2 Gene-expression data

The gene-expression data set was assembled by manually curating bulk profiles from the NCBI Gene Expression Omnibus (GEO) (Barrett et al., 2013). It contains 1,211 samples from six GEO series (GSE10325, GSE32591, GSE38351, GSE39088, GSE45291 and GSE72747) measured on four Affymetrix platforms (GPL96, GPL570, GPL13158 and GPL14663). The sample labels (SLE, Healthy Control or Other/Unknown) were derived from the GEO metadata annotations in the title and characteristics_ch1 fields of each GSM entry. Thirteen genes overlapping the SNP pathways were extracted: *MTHFR*, *TYMS*, *MTR*, *MTRR*, *CYP1A1*, *GSTM1*, *GSTT1*, *GSTP1*, *TLR2*, *TLR4*, *TLR7*, *TLR9* and *STAT4*. The full curated data set is available in the project GitHub repository (https://github.com/catdisk04/AIBD_project).

While re-checking the annotations we found one labelling error. GSE72747 is a lupus nephritis cohort sampled at three time points (T0, T12 and T24), but the ten T0 samples had been labelled Healthy Control. Since all 30 samples in this series come from patients, we relabelled the T0 samples as SLE. After this correction the cohort contains 453 SLE, 189 Healthy Control and 569 Other/Unknown samples (Table 1). The Other/Unknown group includes, for example, monocytes from rheumatoid arthritis patients (GSE38351) and lupus nephritis kidney biopsies (GSE32591), together with samples whose annotation did not clearly indicate SLE or healthy status. These samples were used only to estimate the batch-correction parameters and were never classified.

**Table 1:** Expression batches (GEO series *×* platform). Subjects are distinct individuals inferred from the GEO sample titles.

| Batch | Samples | Healthy | SLE | Other/Unknown | Subjects |
| --- | --- | --- | --- | --- | --- |
| GSE10325_GPL96 | 67 | 28 | 39 | 0 | 25 |
| GSE32591_GPL14663 | 93 | 29 | 0 | 64 | 58 |
| GSE38351_GPL570 | 40 | 36 | 0 | 4 | 15 |
| GSE38351_GPL96 | 34 | 12 | 14 | 8 | 32 |
| GSE39088_GPL570 | 142 | 64 | 78 | 0 | 90 |
| GSE45291_GPL13158 | 805 | 20 | 292 | 493 | 805 |
| GSE72747_GPL570 | 30 | 0 | 30 | 0 | 10 |
| Total | 1211 | 189 | 453 | 569 |  |

#### Repeated measurements

Several series profile the same person more than once. GSE10325 contains CD4^+^ T cells, CD19^+^ B cells and myeloid cells from each donor. GSE39088 and GSE72747 contain three samples per patient. GSE38351 contains monocytes from the same healthy donors under up to five culture conditions, along with technical replicates, and the tubulointerstitial and glomerular biopsies in GSE32591 appear to be paired by donor. We parsed a subject identifier from each sample title; where the title did not identify the person, each sample was treated as a separate individual. The 642 SLE and control samples come from 483 distinct individuals.

## 3 Methodology

### 3.1 Gene-expression data profiling

Before any normalisation or analysis, we first characterised the structure of the raw expression data to understand what we were working with. The data set was classified as bulk gene expression rather than single-cell. This distinction matters because bulk profiles reflect the aggregate transcriptional state of a mixed cell population, and any observed differences between disease and healthy samples represent a combination of true gene regulatory changes and shifts in cell-type composition. Metadata columns (GSE ID, GSM ID, GPL platform and title) were separated from the gene expression columns at the loading stage and excluded from any numerical computation.

### 3.2 Data normalisation and batch correction

Given the multi-study, multi-platform origin of the expression data, normalisation was applied carefully to ensure that biological signal was preserved while technical variability was minimised.

#### Batch definition

Samples measured on different platforms differ technically even within one GEO series, so we defined a batch as series *×* platform. GSE38351 contains samples on both GPL570 and GPL96, which gives seven batches. Disease is spread very unevenly across these batches: two batches contain no SLE samples, one contains no healthy controls, and GSE45291 contributes 292 of the 453 SLE samples but only 20 controls.

#### Scale harmonisation

Per-batch summaries of the values showed three different scales: raw intensities (GSE10325 and GSE38351, with medians in the tens to hundreds and maxima in the thousands), log-scale values (GSE32591 and GSE45291, with medians around 3–10) and values centred near zero (GSE39088 and GSE72747, with medians near zero and negative values). ComBat assumes that batches are on comparable scales, so we harmonised them as follows. Values of *−*1.00 in the raw-intensity series had no within-batch variance and acted as missing-value flags; we set these, and any other non-positive values in those series, to missing. The two raw-intensity series were then transformed as log_2_(*x*) with no pseudocount. The other four series were left on their original scale. The two centred series cannot be returned to the log_2_ scale, and their location difference from the other batches is handled by the mean adjustment in ComBat. The resulting expression values are on a harmonised, log_2_-like scale. These steps act on each sample separately and do not use the disease labels.

#### Gene filtering

ComBat cannot estimate batch parameters for a gene with zero variance or missing values within a batch. A gene was excluded if, in any batch, its standard deviation was zero or undefined, or if it had any missing value. *TLR9* and *GSTM1* met this condition, leaving 11 genes. No samples were removed.

#### ComBat batch correction

Batch effects were corrected with ComBat (Johnson et al., 2007; Leek et al., 2012), an empirical Bayes method that estimates a location (mean) and optionally a scale (variance) adjustment for each gene in each batch. It shrinks these batch parameters toward a prior estimated across genes, which stabilises the estimates for small batches. Disease state was passed as a covariate, with dummy variables for SLE and Other/Unknown and Healthy Control as the reference level, so that ComBat would preserve biological group differences while removing batch effects. We compared two variants: full ComBat (location and scale adjustment) and mean-only ComBat (location adjustment only, variance left unchanged).

We implemented ComBat following the parametric model used in sva and pyComBat (Behdenna et al., 2023), but with the fitting step separated from the correction step, so that batch parameters could be estimated on a training fold and applied to a test fold. Run on the same input as an earlier pyComBat correction of these data, our implementation differed from it by at most 0.051 log_2_ units (mean absolute difference 0.011).

For mean-only ComBat, the corrected value of gene *g* in sample *j* from batch *b* is

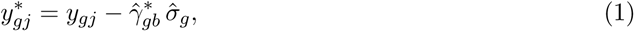

where 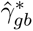 is the shrunken batch location and 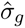 is the pooled residual standard deviation of gene *g*. The disease covariate is needed to estimate 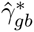, but the correction itself depends only on the batch of the sample. A new, unlabelled sample can therefore be corrected using parameters learned from the training data alone. Full ComBat also rescales each residual around the covariate-predicted mean, which requires the label of the sample, so it cannot be applied to a test sample without using its label. For this reason, and because full ComBat was unstable with only 11 genes (Section 4.1), mean-only ComBat was used as the primary correction and full ComBat is reported as a sensitivity analysis.

### Evaluation of batch correction

Correction was assessed before and after on the 11 retained genes using: (i) the variance explained by batch, computed for each gene as *R*^2^ = *SS*_between_*/SS*_total_ from a one-way ANOVA on batch and averaged across genes; (ii) the batch *R*^2^ of the first two principal components, with PCA run on the gene-standardised matrix; (iii) silhouette scores with Euclidean distance on the gene-standardised data, once with batch labels and once with disease labels; (iv) the same *R*^2^ calculation with disease labels, to check that biological signal was retained; and (v) Cohen’s *d* for SLE versus control, computed per gene with a pooled standard deviation in each batch with at least three SLE and three control samples, together with its spread and sign concordance across batches. To confirm that the disease labels did not drive the correction, we also ran mean-only ComBat without the disease covariate.

### 3.3 Leakage-free evaluation of the expression classifiers

Two sources of leakage were present in the earlier analysis. The normalisation (and, in the first ComBat run, the batch correction) was fitted once on all samples, including the test samples and their labels; and folds were split by sample, so that the same person could contribute, for example, a T-cell profile to training and a B-cell profile to testing. To measure the effect of each, we evaluated the same models under four protocols:

#### P0 (original report)

Globally z-scored 13-gene matrix, original labels, folds split by sample. **P1.** Mean-only ComBat fitted once on all samples with the disease covariate, folds split by sample. **P2.** As P1, but all samples from one individual are kept in the same fold.

#### P3 (leakage-free, primary)

Folds split by individual. Within each training fold, gene filtering, ComBat (fitted on the training SLE and control samples together with the unlabelled Other/Unknown samples), feature standardisation and the classifier are all estimated. The test fold is corrected with Eq. (1) using the training-fold batch parameters and is not used at any point during fitting.

Folds were built with stratified group *k*-fold in scikit-learn (Pedregosa et al., 2011), grouping on subject and stratifying on the combination of batch and label. In every fold, all batches in the test fold were also present in the training fold, and no subject appeared in both (both conditions are checked in the code). Each protocol was run as 10 *×* 5-fold cross-validation with different random splits. Two classifiers were used: L1-penalised logistic regression (L1-LR; *C* = 1) and XGBoost (Chen and Guestrin, 2016) with the same settings as for the SNP data.

#### Confounding checks

Because disease is unevenly distributed across batches, a classifier can score well simply by recognising which study a sample came from. We therefore added three analyses to protocol P3: (a) a batch-only logistic regression whose only input is the one-hot encoded batch label, which shows how much discrimination is available from study membership alone; (b) within-batch ROC-AUC, PR-AUC and MCC, computed from the P3 out-of-fold predictions separately for each batch that contains both classes and combined as a mean weighted by batch size; and (c) leave-one-batch-out (LOBO) validation, in which a whole batch is held out. Because a held-out batch has no ComBat parameters, its gene means were shifted onto the training-set means without using its labels.

#### Model interpretation

The leakage-free XGBoost model was explained with out-of-fold SHAP values (Lundberg and Lee, 2017; Lundberg et al., 2020): in every fold, the model was explained only on the test samples it had not seen, and each sample’s SHAP values were averaged over the 10 repeats. The direction of each gene’s effect was summarised by the Spearman correlation between its expression and its SHAP value. For the L1-LR model we recorded the standardised coefficients of all 50 fold models and report their mean, SD and the fraction of folds with the majority sign. Because a gene can be important to a model only because it helps to tell studies apart, we also computed for each gene a univariate ROC-AUC for SLE over all samples (after global mean-only ComBat) and a within-batch univariate ROC-AUC, averaged over the four batches containing both classes and weighted by batch size. Within-batch AUC uses only the ranking of samples inside a batch, so it is not affected by batch shifts. An AUC above 0.5 means higher expression in SLE.

### 3.4 Genotype analysis

#### Genotype QC

Missingness was calculated for each of the 30 SNPs. Hardy–Weinberg equilibrium (HWE) was then checked in the control group with an exact test (Wigginton et al., 2005), giving a *p*-value per variant. Any variant with HWE *p <* 0.001 was flagged as a possible genotyping error, but kept in the analysis, since the deviation could also reflect a real biological signal.

#### Per-SNP association

For each SNP, a *χ*^2^ test of independence was applied to the 3*×*2 genotype-by-outcome contingency table. Nominal *p*-values were corrected for multiple testing across the 30 variants using the Benjamini–Hochberg (BH) procedure (Benjamini and Hochberg, 1995) at FDR *<* 0.05.

#### Genetic encoding

Three inheritance models were evaluated for every SNP: additive (dose 0/1/2), dominant (any alternate allele *→* 1) and recessive (homozygous alternate only *→* 1). Each encoding was tested alongside age and sex as covariates.

#### Classification

L2-penalised logistic regression (LR; *C* = 1) and XGBoost (300 estimators, maximum depth 3, learning rate 0.03, row and column subsampling 0.8) were evaluated under stratified 10 *×* 5-fold cross-validation. Missing genotypes were imputed with the most frequent value and features were standardised, with both steps fitted inside each training fold. Choosing the best encoding after looking at the cross-validated scores is itself a mild form of selection on the test folds, so we also ran a nested version in which the encoding was chosen by an inner 5-fold cross-validation on the training fold only.

#### Per-SNP effect sizes

To see which genotypes drive each association, we tabulated genotype counts in cases and controls and fitted, for each SNP separately, a logistic regression of SLE on the SNP together with age and sex, under dominant (coded 1 or 2 versus 0), recessive (2 versus 0 or 1) and additive coding. Odds ratios (ORs) are reported with 95% confidence intervals; an OR was not estimated when a cell of the corresponding 2 *×* 2 table was empty. For the carrier comparison we also report Fisher’s exact test, which remains valid when a cell is empty. BH correction was applied across SNPs.

#### Pathway-level models and genotype quality

We grouped the SNPs by the four pathways named by Rupasree et al. (2016): folate and one-carbon metabolism (GCPII, RFC, TYMS 5^′^, TYMS 3^′^, MTHFR, MTR, MTRR A66G), xenobiotic metabolism (CYP1A1 M1, M2 and M4, COMT, GST T1, GST M1), Toll-like receptor signalling (TLR4 299 and 399, two TLR5 variants, TLR7, two TLR9 variants, TIRAP S180L), and STAT4 and other immune or hormonal genes (three STAT4 variants, the TNF*α* CCC haplotype, PTPN22, OPN, two oestrogen receptor variants and IRF5). For each pathway, an LR model with that pathway’s SNPs (dominant coding) plus age and sex was evaluated on the same folds as the main models and compared with age and sex alone. As a check on genotype quality, the additive and dominant models were also refitted without the four SNPs that deviated from HWE in controls (*p <* 0.001).

#### Interpretability

SHAP (SHapley Additive exPlanations) values (Lundberg and Lee, 2017; Lundberg et al., 2020) were computed out of fold: in every training fold, the XGBoost model (additive coding plus age and sex) was explained on the held-out subjects only, and each subject’s SHAP values were averaged over the 10 repeats. The direction of each feature’s effect was summarised by the Spearman correlation between the feature value and its SHAP value.

### 3.5 Performance metrics and model comparison

For every test fold we computed ROC-AUC, PR-AUC (average precision), MCC, F1, balanced accuracy, accuracy and the Brier score, using a probability threshold of 0.5 for the threshold-dependent metrics. PR-AUC is reported because the expression data are imbalanced (70.6% SLE), and in that setting ROC-AUC can give an overly favourable picture (Saito and Rehmsmeier, 2015). The chance level of PR-AUC equals the prevalence of the positive class: 0.706 for the expression data and 0.442 for the genotype data. MCC is reported because, unlike accuracy and F1, it is high only when the classifier performs well on both classes (Chicco and Jurman, 2020). We report the mean *±* SD over all 50 folds. Two models evaluated on the same folds were compared with the corrected resampled *t*-test of Nadeau and Bengio (2003), which accounts for the overlap between training sets in repeated cross-validation.

### 3.6 Permutation checks

If an evaluation pipeline leaks test information, it can score above chance even when the labels carry no information. We therefore re-ran the complete pipelines with shuffled labels, 100 times each, on the first cross-validation split. For the SNP data (dominant LR), labels were shuffled across subjects. For the expression data (P3, both classifiers), labels were shuffled in two ways: across all samples, with the shuffled labels also passed to ComBat, and only within each batch, which keeps the batch–label confounding but removes any link between gene expression and disease inside a batch. The permutation *p*-value is (1 + #*{*null *≥* observed*}*)*/*(1 + 100), so the smallest possible value is 0.0099.

### 3.7 Differential expression

Following Nygaard et al. (2016), differential expression was tested on the harmonised (uncorrected) data with batch included in the model, rather than on the ComBat output. Testing on data that were corrected using the disease covariate can overstate the significance of disease effects. For each gene we fitted expression *∼* SLE +batch by ordinary least squares on the SLE and control samples, with standard errors clustered by subject, and applied BH-FDR correction across the 11 genes.

## 4 Results

### 4.1 Batch correction

Before correction, batch explained 91.9% of the variance of the average gene, and the first principal component separated the two centred series from the rest (Fig. 1A; PC1 batch *R*^2^ = 0.970). Mean-only ComBat reduced the mean batch *R*^2^ to 0.131, the PC1 batch *R*^2^ to 0.203 and the batch silhouette from 0.404 to 0.056, while the disease silhouette increased from 0.141 to 0.235 (Table 2; Fig. 1B and D). Full ComBat removed more batch variance (mean batch *R*^2^ 0.060) but distorted within-dataset effects. It roughly doubled the spread of Cohen’s *d* across datasets (from 0.554 to 1.144) and produced an implausible SLE-versus-control effect of *d* = *−*13.2 for *CYP1A1* in GSE45291, which we attribute to unstable scale estimates from only 11 genes. Mean-only correction leaves within-batch differences unchanged by construction, which is why its Cohen’s *d* values match the uncorrected ones.

**Figure 1:**
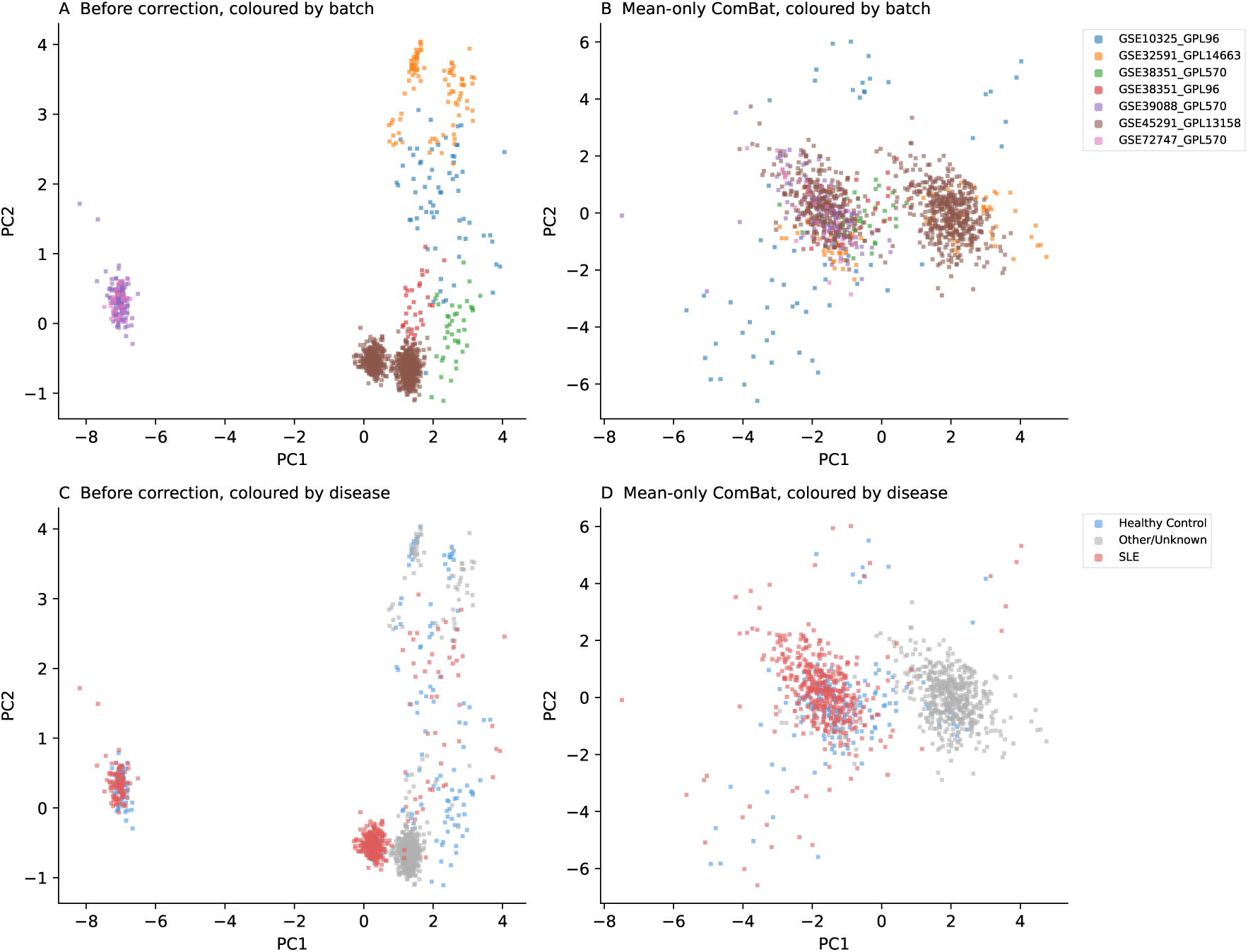
PCA of the 11-gene expression matrix before and after mean-only ComBat. (A, B) Coloured by batch; (C, D) coloured by disease state. Before correction the samples cluster by study and platform; after correction the batches overlap and the SLE samples separate from the Other/Unknown samples.

**Table 2:** Batch-correction diagnostics on the 11 retained genes. Cohen’s *d* statistics use the four batches with at least three SLE and three control samples.

| Metric | Before | Full ComBat | Mean-only ComBat | Mean-only, no covariate |
| --- | --- | --- | --- | --- |
| Mean batch $R^2$ (genes) | 0.919 | 0.060 | 0.131 | 0.095 |
| Mean disease $R^2$ (genes) | 0.176 | 0.287 | 0.265 | 0.192 |
| PC1 batch $R^2$ | 0.970 | 0.143 | 0.203 | 0.017 |
| PC2 batch $R^2$ | 0.938 | 0.014 | 0.043 | 0.117 |
| Silhouette (batch) | 0.404 | −0.113 | 0.056 | 0.040 |
| Silhouette (disease) | 0.141 | 0.105 | 0.235 | 0.201 |
| SD of Cohen’s $d$ across batches | 0.554 | 1.144 | 0.554 | 0.554 |

The disease covariate also raises the apparent disease signal. Without it, mean-only ComBat gave a mean disease *R*^2^ of 0.192, compared with 0.265 when the covariate was included. This is the effect described by Nygaard et al. (2016), and for this reason no statistical test in this study was run on the corrected matrix.

### 4.2 SNP QC and variant association

Overall missingness was low (mean 0.2%, maximum 4.7% for CYP1A1M1). HWE testing in controls flagged four variants at *p <* 0.001, two of them with extreme deviations: *MTRR* A66G (*p* = 1.84 *×* 10^−12^) and *TLR5* rs2072493 (*p* = 1.52 *×* 10^−10^) (Fig. 2A). These deviations may reflect local genotyping difficulty rather than true biological enrichment. Per-SNP *χ*^2^ association testing identified four variants passing FDR *<* 0.05 (Fig. 2B): *MTRR* A66G (*q* = 0.0002), CYP1A1M2 (*q* = 0.014), *COMT* (*q* = 0.032) and CYP1A1M4 (*q* = 0.044). Seven additional SNPs reached nominal significance (*p <* 0.05), including *MTHFR* (*p* = 0.034), *OPN* (*p* = 0.032) and the *TNF α* CCC haplotype (*p* = 0.039). The strongest association, at *MTRR* A66G, is also one of the two most extreme HWE deviations, so the genotype pattern behind it needs a closer look.

**Figure 2:**
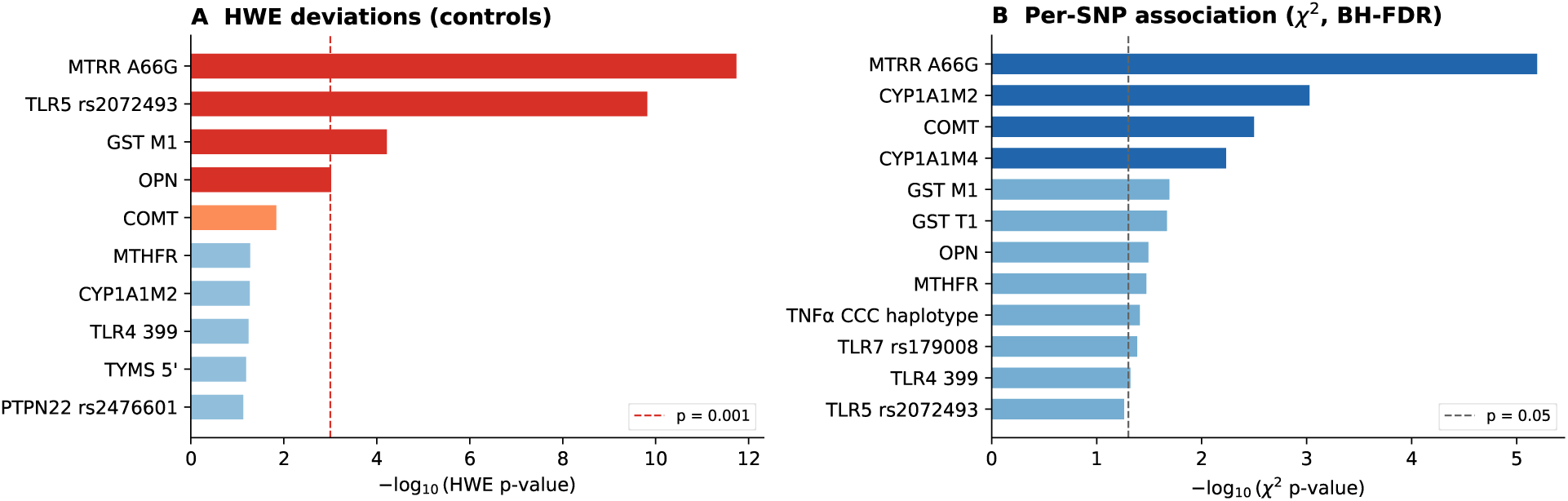
SNP quality control and association. (A) HWE *p*-values in controls (*−* log_10_ scale); dashed line at *p* = 0.001. (B) Per-SNP *χ*^2^ association strength; dark blue bars indicate BH-FDR *q <* 0.05.

### 4.3 Genotype patterns behind the SNP associations

Table 3 lists the genotype counts and odds ratios for the 11 SNPs with nominal *χ*^2^ association. The signals fall into three groups.

**Table 3:** Genotype counts and effect sizes for the 11 SNPs with nominal *χ*^2^ association (*p <* 0.05). Counts are for genotypes coded 0/1/2 in the source data. ORs (95% CI) are from logistic regression on one SNP plus age and sex; dominant: coded 1 or 2 vs. 0; recessive: 2 vs. 0 or 1. Fisher: exact test of carriers vs. non-carriers. n.e.: not estimable (empty cell).

| SNP | Pathway | SLE 0/1/2 | Control 0/1/2 | HWE $p$ | $\chi^2 q$ | Dominant OR | Fisher $p$ | Recessive OR |
| --- | --- | --- | --- | --- | --- | --- | --- | --- |
| *MTRR* A66G | Folate | 21/117/50 | 3/186/48 | $1.8 \times 10^{-12}$ | $1.9 \times 10^{-4}$ | 0.12 (0.04–0.42) | $8.9 \times 10^{-6}$ | 1.50 (0.94–2.40) |
| *CYP1A1* M2 | Xenobiotic | 134/41/13 | 163/72/2 | 0.054 | 0.014 | 0.78 (0.51–1.21) | 0.60 | 7.78 (1.71–35.43) |
| *COMT* | Xenobiotic | 69/78/41 | 79/131/27 | 0.015 | 0.032 | 0.89 (0.59–1.34) | 0.48 | 2.31 (1.34–3.99) |
| *CYP1A1* M4 | Xenobiotic | 180/7/1 | 237/0/0 | 1.00 | 0.044 | n.e. | 0.0013 | n.e. |
| *GST M1* | Xenobiotic | 139/49/0 | 149/88/0 | $6.1 \times 10^{-5}$ | 0.11 | 0.65 (0.42–1.00) | 0.016 | n.e. |
| *GST T1* | Xenobiotic | 129/59/0 | 187/50/0 | 0.086 | 0.11 | 1.71 (1.09–2.68) | 0.019 | n.e. |
| *OPN* | Immune | 78/74/36 | 128/77/32 | $9.6 \times 10^{-4}$ | 0.12 | 1.61 (1.08–2.39) | 0.011 | 1.59 (0.93–2.72) |
| *MTHFR* | Folate | 160/28/0 | 181/56/0 | 0.053 | 0.12 | 0.57 (0.34–0.96) | 0.027 | n.e. |
| *TNFα* CCC | Immune | 172/16/0 | 229/8/0 | 1.00 | 0.12 | 2.53 (1.04–6.14) | 0.033 | n.e. |
| *TLR7* rs179008 | TLR | 179/7/2 | 211/24/2 | 0.19 | 0.12 | 0.38 (0.17–0.85) | 0.022 | 1.17 (0.16–8.57) |
| *TLR4* 399 | TLR | 124/58/6 | 172/64/1 | 0.058 | 0.13 | 1.34 (0.87–2.05) | 0.17 | 7.78 (0.91–66.20) |

First, the association at *MTRR* A66G does not follow an allele-dose pattern. The genotype coded 0 was carried by 11.2% of cases but only 1.3% of controls, heterozygotes made up 78.5% of controls compared with 62.2% of cases, and the additive model showed no trend (OR 0.93, *p* = 0.71). An excess of heterozygotes in controls is the same pattern that produces the extreme HWE deviation (*p* = 1.8 *×* 10^−12^), and it is a typical signature of genotyping error (Hosking et al., 2004). This association should therefore be treated with caution until it is replicated with an independent genotyping method.

Second, the xenobiotic-metabolism variants show effects concentrated in homozygotes or rare carriers. The genotype coded 2 at *CYP1A1* M2 was found in 6.9% of cases and 0.8% of controls (recessive OR 7.78, 95% CI 1.71–35.43), and at *COMT* in 21.8% of cases and 11.4% of controls (recessive OR 2.31, 1.34–3.99). *CYP1A1* M4 carriers were rare but occurred only among cases (8 of 188 versus 0 of 237; Fisher *p* = 0.0013, *q* = 0.020). Carriers at *GST T1* were more frequent among cases (31.4% versus 21.1%; OR 1.71, 1.09–2.68), and carriers at *GST M1* less frequent (26.1% versus 37.1%; OR 0.65, 0.42–1.00). These recessive and rare-variant patterns explain why *CYP1A1* M2 and *COMT* reached FDR significance in the three-genotype *χ*^2^ test but not under dominant coding.

Third, several immune and folate variants showed modest carrier effects at nominal significance only: the *TNF α* CCC haplotype (8.5% versus 3.4%; OR 2.53, 1.04–6.14), *OPN* (58.5% versus 46.0%; OR 1.61, 1.08–2.39), *MTHFR* (14.9% versus 23.6%; OR 0.57, 0.34–0.96) and *TLR7* rs179008 (4.8% versus 11.0%; OR 0.38, 0.17–0.85). Apart from *MTRR* A66G and *CYP1A1* M4, none of the carrier comparisons survived FDR correction (Fisher *q ≥* 0.108).

### 4.4 SNP classification performance

Table 5 summarises the cross-validated performance of the SNP models. Age and sex alone gave ROC-AUC 0.633 and PR-AUC 0.558 (chance level 0.442), confirming that these covariates carry limited discriminative power. Adding the SNPs under a dominant encoding raised these to 0.699 and 0.668, and MCC rose from 0.176 to 0.309. The dominant encoding outperformed the additive encoding on ROC-AUC (difference 0.044, corrected *p* = 0.023) and PR-AUC (difference 0.046, *p* = 0.038); the MCC difference (0.097, *p* = 0.080) was not significant. The recessive encoding performed worst. When the encoding was chosen inside each training fold, the dominant encoding was picked in 48 of the 50 folds and performance was essentially unchanged (ROC-AUC 0.696, PR-AUC 0.664), so the support for H1 does not depend on choosing the encoding after seeing the test folds. XGBoost with additive encoding (ROC-AUC 0.690) did not differ significantly from LR with the same features (ROC-AUC *p* = 0.120; PR-AUC *p* = 0.410), consistent with H3.

Table 4 breaks the SNP signal down by pathway and genotype quality. No single pathway came close to the full dominant model. The folate and xenobiotic SNP sets gave the largest gains over age and sex (ROC-AUC 0.653 and 0.647), significant for PR-AUC (*p* = 0.020 and *p* = 0.045) but not for ROC-AUC (*p* = 0.40 and *p* = 0.57), while the TLR and STAT4/immune sets added nothing (0.613 and 0.631). When the four SNPs that fail HWE in controls (*MTRR* A66G, *TLR5* rs2072493, *GST M1* and *OPN*) were removed, the dominant model fell to ROC-AUC 0.656 *±* 0.053, PR-AUC 0.605 *±* 0.062 and MCC 0.231 *±* 0.104. It was then no better than the additive model (ROC-AUC difference 0.008, *p* = 0.65) and not significantly better than age and sex alone (ROC-AUC *p* = 0.39, PR-AUC *p* = 0.16). Much of the SNP signal, and the advantage of dominant coding, therefore rests on SNPs whose genotypes may be unreliable.

**Table 4:** SNP models by pathway and genotype quality (LR, dominant coding unless stated, all with age and sex; 10 *×* 5-fold CV, mean *±* SD). The last column is the corrected resampled *t*-test of ROC-AUC against age and sex alone.

| SNP set | ROC-AUC | PR-AUC | MCC | $p$ vs. age + sex |
| --- | --- | --- | --- | --- |
| None (age + sex only) | $0.633 \pm 0.056$ | $0.558 \pm 0.055$ | $0.176 \pm 0.076$ | – |
| All 30 SNPs | $0.699 \pm 0.052$ | $0.668 \pm 0.063$ | $0.309 \pm 0.101$ | 0.024 |
| Folate / one-carbon (7) | $0.653 \pm 0.060$ | $0.617 \pm 0.060$ | $0.197 \pm 0.112$ | 0.40 |
| Xenobiotic metabolism (6) | $0.647 \pm 0.044$ | $0.620 \pm 0.058$ | $0.185 \pm 0.101$ | 0.57 |
| Toll-like receptor (8) | $0.613 \pm 0.050$ | $0.553 \pm 0.049$ | $0.125 \pm 0.095$ | 0.42 |
| STAT4 and other immune (9) | $0.631 \pm 0.062$ | $0.570 \pm 0.068$ | $0.196 \pm 0.113$ | 0.92 |
| 26 HWE-passing SNPs, additive | $0.648 \pm 0.049$ | $0.599 \pm 0.053$ | $0.186 \pm 0.094$ | – |
| 26 HWE-passing SNPs, dominant | $0.656 \pm 0.053$ | $0.605 \pm 0.062$ | $0.231 \pm 0.104$ | 0.39 |

**Table 5:**
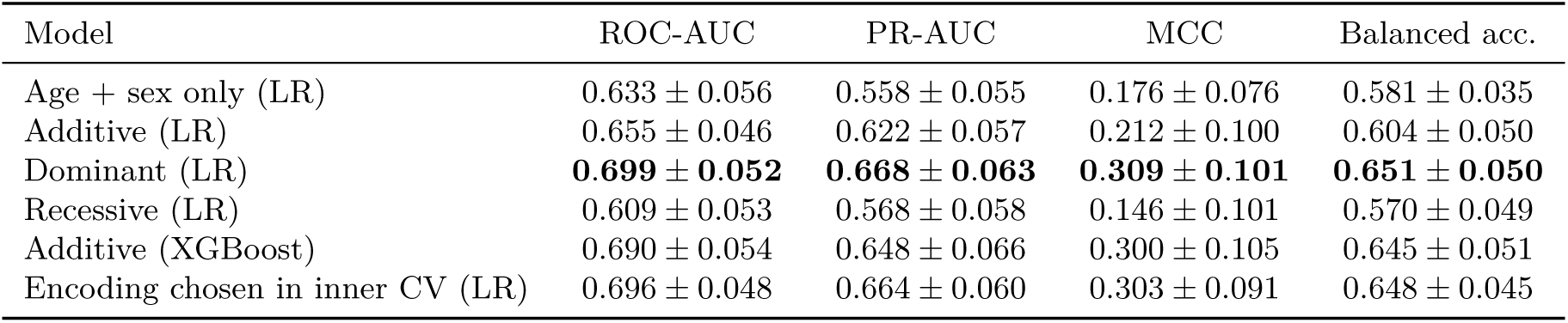
SNP classification, 10 *×* 5-fold stratified cross-validation (mean *±* SD over 50 folds). All models include age and sex. The PR-AUC chance level is 0.442.

| Model | ROC-AUC | PR-AUC | MCC | Balanced acc. |
| --- | --- | --- | --- | --- |
| Age + sex only (LR) | $0.633 \pm 0.056$ | $0.558 \pm 0.055$ | $0.176 \pm 0.076$ | $0.581 \pm 0.035$ |
| Additive (LR) | $0.655 \pm 0.046$ | $0.622 \pm 0.057$ | $0.212 \pm 0.100$ | $0.604 \pm 0.050$ |
| Dominant (LR) | **$0.699 \pm 0.052$** | **$0.668 \pm 0.063$** | **$0.309 \pm 0.101$** | **$0.651 \pm 0.050$** |
| Recessive (LR) | $0.609 \pm 0.053$ | $0.568 \pm 0.058$ | $0.146 \pm 0.101$ | $0.570 \pm 0.049$ |
| Additive (XGBoost) | $0.690 \pm 0.054$ | $0.648 \pm 0.066$ | $0.300 \pm 0.105$ | $0.645 \pm 0.051$ |
| Encoding chosen in inner CV (LR) | $0.696 \pm 0.048$ | $0.664 \pm 0.060$ | $0.303 \pm 0.091$ | $0.648 \pm 0.045$ |

Out-of-fold SHAP values for the XGBoost model (Fig. 6A) ranked age, *OPN*, *MTRR* A66G, *COMT* and *GST T1* as the top five contributors. Younger age pushed predictions toward SLE, reflecting the younger case group (mean 27.4 versus 31.4 years in controls) rather than a genetic effect. For *MTRR* A66G, the largest positive SHAP values came from the rare genotype coded 0, matching the genotype counts above. Three of the six highest-ranked features (*OPN*, *MTRR* A66G and *GST M1*) are SNPs that fail HWE in controls.

### 4.5 Expression classification and data leakage

Table 6 and Fig. 3 show the expression classifiers under the four protocols. Protocol P0 reproduces the result of the original report (ROC-AUC 0.807). Replacing the z-scores with ComBat-corrected values (P1) improved L1-LR to ROC-AUC 0.832 and MCC 0.475. Moving to subject-grouped folds (P2) and then fitting ComBat inside each fold (P3) changed the mean of every metric by at most 0.015, and the leaky protocols scored slightly lower rather than higher. The leakage-free L1-LR reached ROC-AUC 0.837 *±* 0.033, PR-AUC 0.920 *±* 0.023 (chance level 0.706) and MCC 0.485 *±* 0.073. XGBoost performed better on every metric (ROC-AUC 0.915, PR-AUC 0.963, MCC 0.640), and its advantage over L1-LR was significant (corrected *p <* 10^−4^ for ROC-AUC, *p* = 0.0005 for PR-AUC and *p* = 0.0003 for MCC). With these data, therefore, neither the global ComBat fit nor the repeated subjects noticeably inflated the cross-validated estimates.

**Figure 3:**
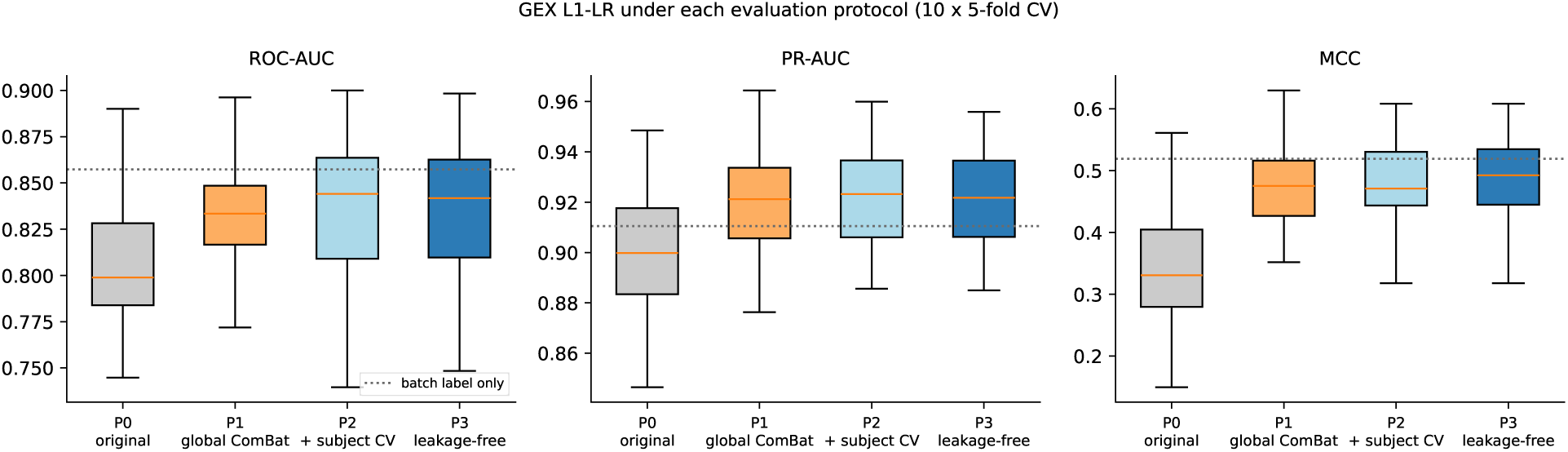
Performance of the expression L1-LR classifier under the four evaluation protocols (50 folds each). The dotted line shows the batch-label-only model under the leakage-free protocol.

**Table 6:**
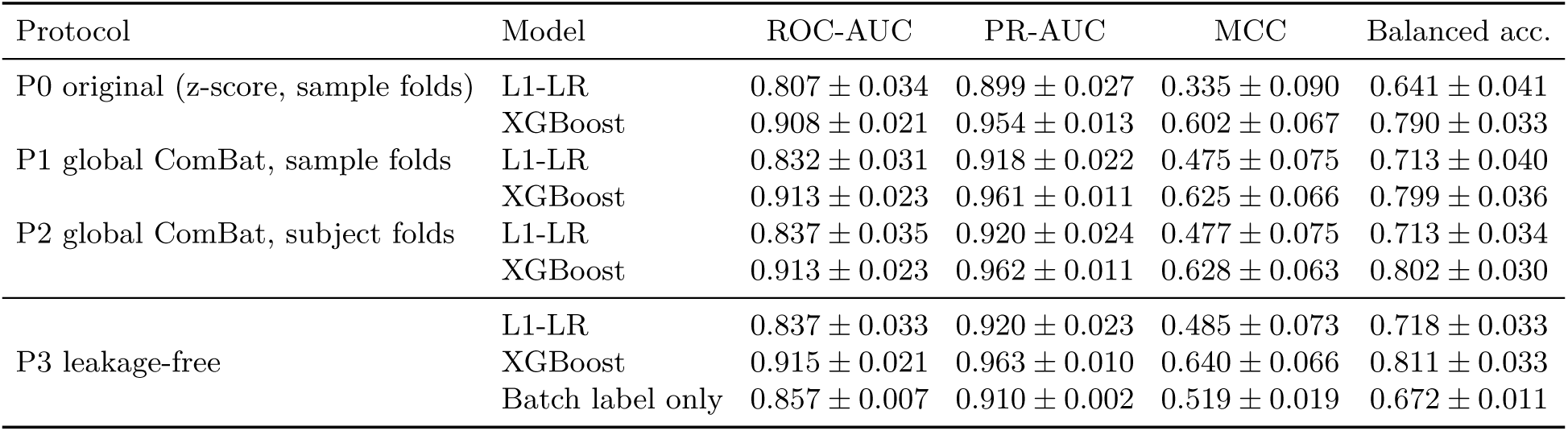
Expression classification (SLE vs. control; 642 samples from 483 individuals), 10 *×* 5-fold cross-validation (mean *±* SD over 50 folds). P0 uses the labels of the original report. The PR-AUC chance level is 0.706.

### 4.6 Study membership explains much of the pooled performance

The confounding checks show that the pooled estimates in Table 6 overstate how well these genes separate SLE from controls. A logistic regression given only the batch label, with no expression data at all, reached ROC-AUC 0.857 and MCC 0.519, higher than the expression L1-LR. This follows from the batch composition in Table 1: a sample from GSE45291 or GSE72747 is almost always SLE, and a sample from GSE32591 or GSE38351_GPL570 is always a control.

Within individual batches, where study membership carries no information, discrimination was lower (Table 7). The within-batch ROC-AUC, weighted by batch size, was 0.743 *±* 0.006 for L1-LR and 0.835 *±* 0.006 for XGBoost, compared with 0.486 for the batch-only model. Most of the signal came from the two whole-blood cohorts, GSE39088 (ROC-AUC 0.784 for L1-LR and 0.855 for XGBoost) and GSE45291 (0.771 and 0.898), and the signal was weak in the sorted-cell and monocyte series GSE10325 and GSE38351_GPL96 (0.576–0.621). In GSE45291, which is 93.6% SLE, L1-LR has a high PR-AUC (0.981) but an MCC close to zero (0.009): at the 0.5 threshold it labels almost every sample as SLE (95.2% of samples in the first cross-validation repeat, including 18 of the 20 controls). ROC-AUC and PR-AUC alone would not have shown this failure, whereas MCC does. When whole batches were held out, ROC-AUC fell to 0.553–0.684 for L1-LR and 0.463–0.741 for XGBoost, with MCC between *−*0.181 and 0.153 (Supplementary Table S1). Figure 4 compares the out-of-fold ROC and precision–recall curves of the two modalities.

**Figure 4:**
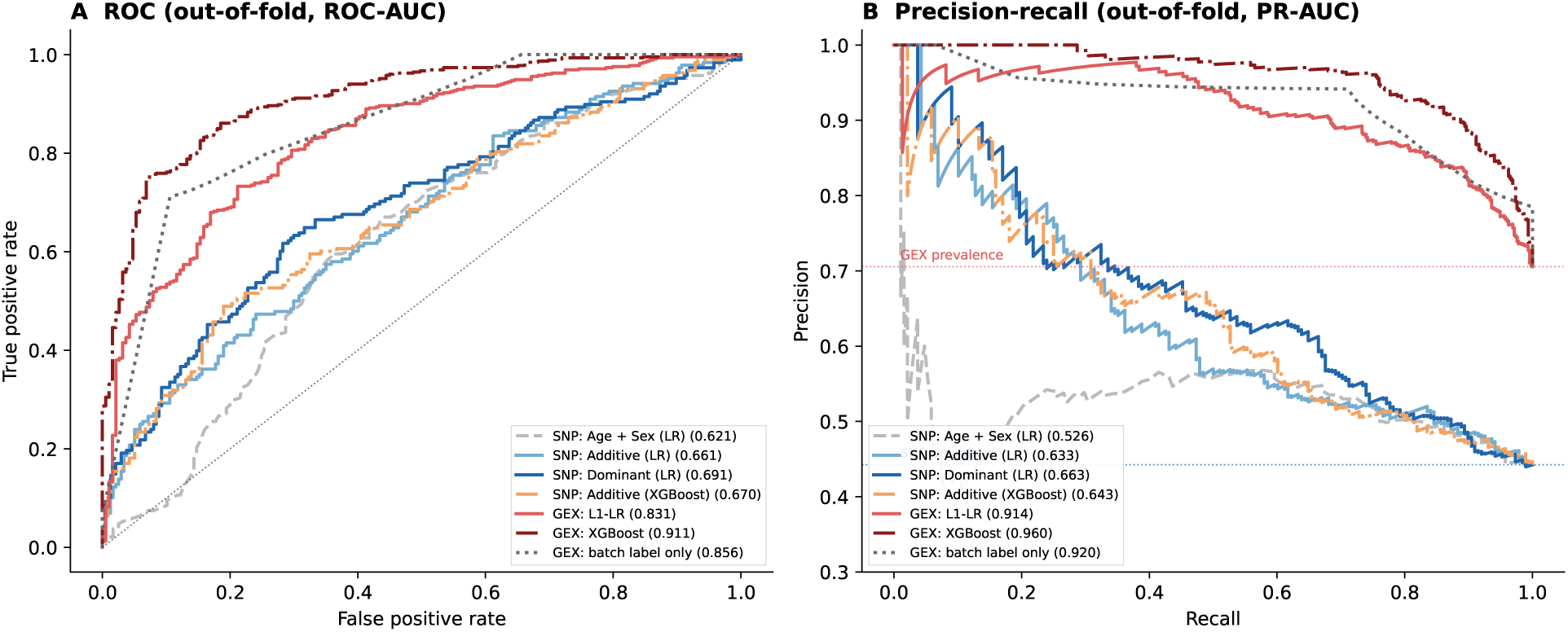
Out-of-fold ROC (A) and precision–recall (B) curves for the SNP models and the leakage-free (P3) expression models, from the first cross-validation repeat (so the values in the legend differ slightly from the 50-fold means in Tables 5 and 6). Dotted horizontal lines in (B) mark the prevalence, which is the chance level of PR-AUC, for each data set. The SNP and expression curves come from different cohorts.

**Table 7:** Within-batch discrimination of the leakage-free (P3) out-of-fold predictions, averaged over the 10 repeats. Only batches containing both classes are shown.

| Batch | $n$ | SLE (%) | L1-LR | | | XGBoost | | |
| --- | --- | --- | --- | --- | --- | --- | --- | --- |
|  |  |  | ROC | PR | MCC | ROC | PR | MCC |
| GSE10325_GPL96 | 67 | 58.2 | 0.588 | 0.724 | 0.066 | 0.580 | 0.656 | 0.166 |
| GSE38351_GPL96 | 26 | 53.8 | 0.576 | 0.651 | 0.150 | 0.621 | 0.607 | 0.247 |
| GSE39088_GPL570 | 142 | 54.9 | 0.784 | 0.836 | 0.398 | 0.855 | 0.899 | 0.535 |
| GSE45291_GPL13158 | 312 | 93.6 | 0.771 | 0.981 | 0.009 | 0.898 | 0.991 | 0.475 |
| Weighted mean | 547 |  | 0.743 | 0.896 | 0.124 | 0.835 | 0.908 | 0.442 |

### 4.7 Differential expression

After adjusting for batch, 4 of the 11 genes were differentially expressed at FDR *<* 0.05 (Fig. 5; Supplementary Table S2): *MTR* (log_2_FC = *−*0.32, *q* = 6 *×* 10^−5^), *TYMS* (+0.34, *q* = 2 *×* 10^−4^), *STAT4* (*−*0.24, *q* = 0.003) and *GSTP1* (*−*0.18, *q* = 0.012). *TLR2* was nominally up-regulated (+0.18, *p* = 0.028, *q* = 0.062). These results differ from our earlier, unadjusted Mann–Whitney analysis on the pooled z-scores, which reported 9 of 13 genes as significant, a large up-regulation of *TLR2* (median log_2_FC = +2.39), down-regulation of *MTRR* and up-regulation of *STAT4*. Those effects largely reflected which studies contributed cases and which contributed controls. Of the earlier findings, only the down-regulation of *MTR* is still supported.

**Figure 5:**
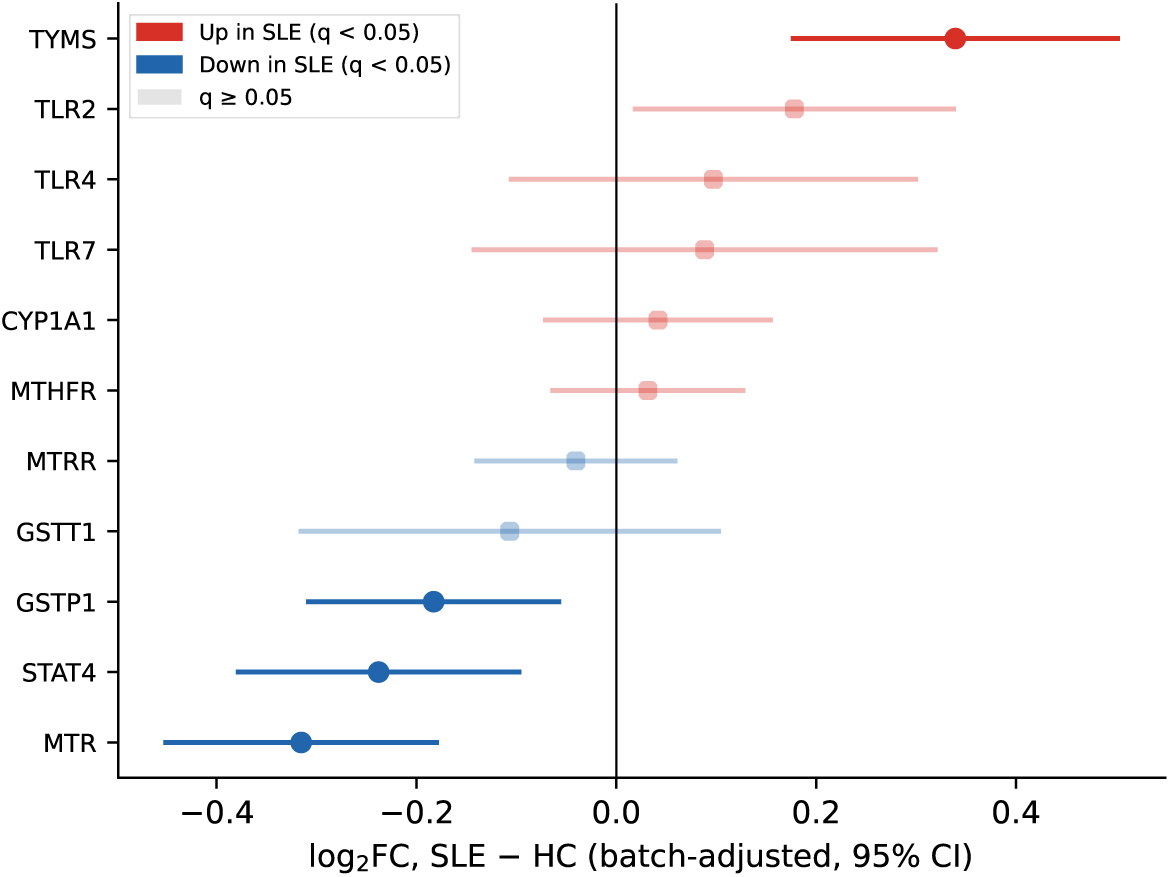
Batch-adjusted differential expression (SLE *−* control) with 95% confidence intervals, from OLS with batch fixed effects and subject-clustered standard errors. Faded points are not significant at BH-FDR *q <* 0.05.

**Figure 6:**
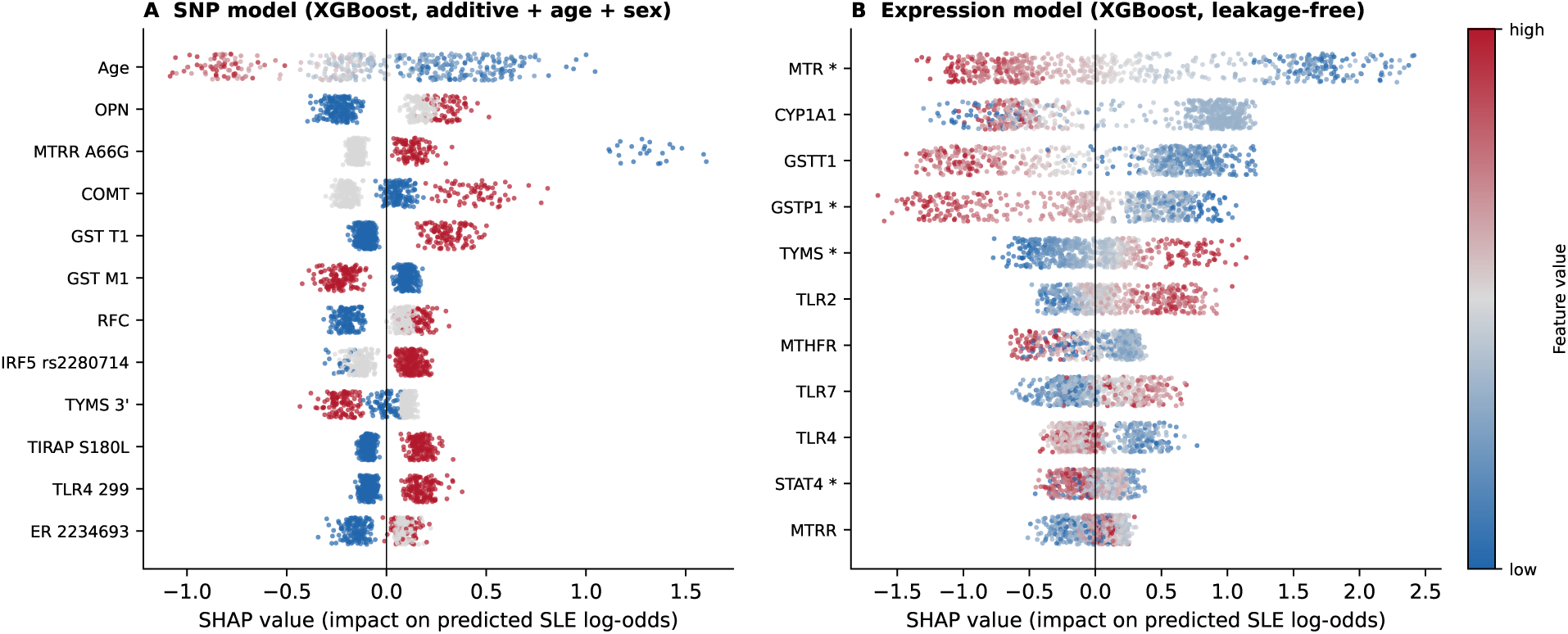
Out-of-fold SHAP values. (A) SNP model (XGBoost, additive coding plus age and sex), 12 highest-ranked features. (B) Leakage-free expression model (XGBoost). Each point is one subject or sample, coloured by its feature value (genotypes 0/1/2 map to low/middle/high). Positive SHAP values push the prediction toward SLE. Asterisks in (B) mark genes differentially expressed after batch adjustment (BH *q <* 0.05).

### 4.8 What the expression models rely on

Figure 6B and Table 8 summarise what the leakage-free expression models used. *MTR* was the most important feature for XGBoost (mean *|*SHAP*|* 0.866), with lower expression pushing predictions toward SLE (Spearman *ρ* = *−*0.96 between expression and SHAP value). It also had the largest L1-LR coefficient (*−*1.386*±*0.125, same sign in all 50 folds), was lower in SLE within batches (within-batch AUC 0.363) and was the most significant gene after batch adjustment. *GSTP1*, *TYMS* and *STAT4* likewise kept the same direction within batches (within-batch AUC 0.318, 0.675 and 0.342) as in the batch-adjusted differential expression analysis, although the L1-LR coefficient of *STAT4* was positive once the other genes were in the model.

**Table 8:** What the leakage-free expression models rely on. *|*SHAP*|*: mean absolute out-of-fold SHAP value (XGBoost); *ρ*: Spearman correlation between expression and SHAP value (negative: lower expression pushes toward SLE). L1-LR: standardised coefficient (mean *±* SD over 50 folds) and fraction of folds with the majority sign. AUC: univariate ROC-AUC for SLE over pooled samples and within batches (below 0.5: lower in SLE). DE: batch-adjusted log_2_FC and BH *q*.

| Gene | XGBoost SHAP |  | L1-LR |  | Univariate AUC |  | DE |  |
| --- | --- | --- | --- | --- | --- | --- | --- | --- |
| | \|SHAP\| | $\rho$ | Coefficient | Sign agreement | Pooled | Within batch | $\log_2$FC | $q$ |
| *MTR* | 0.866 | -0.96 | $-1.386 \pm 0.125$ | 1.00 | 0.237 | 0.363 | -0.32 | $5.6 \times 10^{-5}$ |
| *CYP1A1* | 0.761 | -0.31 | $-0.153 \pm 0.060$ | 0.98 | 0.432 | 0.514 | +0.04 | 0.51 |
| *GSTT1* | 0.705 | -0.81 | $-0.665 \pm 0.096$ | 1.00 | 0.271 | 0.430 | -0.11 | 0.51 |
| *GSTP1* | 0.548 | -0.89 | $-0.553 \pm 0.079$ | 1.00 | 0.399 | 0.318 | -0.18 | 0.012 |
| *TYMS* | 0.290 | +0.97 | $+0.507 \pm 0.115$ | 1.00 | 0.522 | 0.675 | +0.34 | $2.3 \times 10^{-4}$ |
| *TLR2* | 0.249 | +0.89 | $+0.615 \pm 0.152$ | 1.00 | 0.671 | 0.550 | +0.18 | 0.062 |
| *MTHFR* | 0.233 | -0.35 | $-0.054 \pm 0.065$ | 0.78 | 0.390 | 0.635 | +0.03 | 0.51 |
| *TLR7* | 0.232 | +0.76 | $-0.020 \pm 0.061$ | 0.52 | 0.403 | 0.603 | +0.09 | 0.51 |
| *TLR4* | 0.201 | -0.66 | $-0.240 \pm 0.165$ | 0.94 | 0.522 | 0.494 | +0.10 | 0.51 |
| *STAT4* | 0.153 | -0.39 | $+0.130 \pm 0.067$ | 0.96 | 0.497 | 0.342 | -0.24 | 0.0033 |
| *MTRR* | 0.140 | +0.41 | $+0.274 \pm 0.085$ | 1.00 | 0.375 | 0.580 | -0.04 | 0.51 |

The second- and third-ranked genes for XGBoost, *CYP1A1* and *GSTT1*, behaved differently. Both carried little signal inside individual batches (within-batch AUC 0.514 and 0.430) and neither was differentially expressed after batch adjustment (*q* = 0.51), even though *GSTT1* looked strongly lower in SLE when all samples were pooled (pooled AUC 0.271). Their high SHAP values therefore most likely reflect differences between studies that XGBoost can exploit, in line with the batch-only baseline. The same pattern explains the earlier *TLR2* result: pooled AUC 0.671 but within-batch AUC 0.550. Two genes were unstable in the linear model: the sign of the *TLR7* coefficient agreed in only 52% of folds, and that of *MTHFR* in 78%.

### 4.9 Permutation checks

With labels shuffled across all samples, the leakage-free pipelines scored at chance: mean pooled ROC-AUC 0.489 for L1-LR and 0.498 for XGBoost (95th percentiles 0.550 and 0.555), and 0.495 for the SNP model (95th percentile 0.555). The observed values on the same split (0.831, 0.911 and 0.691) exceeded every permuted value (*p* = 0.0099; Supplementary Fig. S1), which shows that the pipeline itself does not leak label information. Shuffling labels only within batches gave a different picture. Under this null the pooled ROC-AUC was 0.733 for L1-LR and 0.811 for XGBoost, purely from batch–label confounding, while the within-batch ROC-AUC stayed at chance (0.489 and 0.485). The observed within-batch ROC-AUC (0.739 and 0.829) was above all permuted values, so the expression models do capture disease-related signal inside studies, but a large share of their pooled performance would be obtained even without any such signal.

## 5 Discussion and Conclusion

### Hypotheses

H1 is only partly supported. The dominant encoding improved ROC-AUC and PR-AUC over the additive encoding, the gain held up under a corrected paired test, and the dominant encoding was the one selected inside the training folds. However, the advantage disappeared when the four SNPs that fail HWE were removed, and much of it came from *MTRR* A66G, whose non-additive genotype pattern is easy to capture with dominant coding. Because the coded allele is not always the minor allele, “dominant” here describes the coding rather than a biological mode of inheritance. H2 is not supported as originally framed. *MTRR* A66G gave the smallest *p*-value, but its association is driven by a heterozygote excess in controls, and the genotype enriched in cases is the one coded 0. If the code counts copies of the 66G allele, as the SNP name suggests, this is the AA genotype, whereas a folate-deficiency mechanism would predict enrichment of the 66G allele, which lowers the efficiency of methionine synthase reactivation (Olteanu et al., 2002). The folate SNP set as a whole added about as much to age and sex as the xenobiotic set did. H3 is supported for the genotype data, where XGBoost did not significantly outperform logistic regression; this is consistent with the limited sample size (*n* = 425), since tree ensembles typically require larger data sets to outperform regularised linear models on structured tabular data. On the larger and more heterogeneous expression data XGBoost was clearly better, both pooled and within batches, although part of its advantage comes from genes that mainly separate studies. H4 is supported only with a qualification. Pooled expression performance (ROC-AUC 0.837–0.915) is well above that of the genotype models (0.699), but a large part of it comes from study membership. Within studies, L1-LR (0.743) is only modestly above the best SNP model, while XGBoost (0.835) remains clearly ahead. The two modalities were also measured in different individuals, so this comparison is indirect.

### Why these SNPs predict SLE

The SNP signal is spread over many small effects rather than concentrated in one pathway, and the most credible part of it lies in xenobiotic metabolism. CYP1A1 is a phase I enzyme that converts polycyclic aromatic hydrocarbons, such as those in tobacco smoke, into reactive intermediates, and the glutathione S-transferases (GST T1 and GST M1) conjugate such intermediates for excretion. Variation in this pathway could change how strongly environmental exposures act on the immune system, which fits with smoking being a risk factor for SLE (Costenbader et al., 2004). COMT methylates catechols, including catechol oestrogens, using *S*-adenosylmethionine (SAM) as the methyl donor, so it links hormone metabolism, which is relevant to the strong female bias of SLE, with the supply of methyl groups. The folate pathway feeds the same methyl supply: MTHFR produces 5-methyltetrahydrofolate, which MTR uses to remethylate homocysteine to methionine, the precursor of SAM, and MTRR keeps MTR active. Reduced DNA methylation in T cells is a long-recognised feature of SLE (Richardson et al., 1990; Javierre et al., 2010). On the expression side, lower *MTR* was the most robust finding in every analysis, and *TYMS*, which uses folate for thymidylate synthesis, was higher in SLE. One possible reading is that folate is directed toward nucleotide synthesis in activated immune cells rather than toward methylation, but our data cannot test this directly. In the genotype data, the folate signal comes mostly from *MTRR* A66G, whose genotype quality is doubtful, and from a modest *MTHFR* carrier effect. The immune variants contributed little as a group; the *TNF α* CCC haplotype and *OPN* showed nominal carrier effects, and *OPN* also fails HWE. Taken together, the SNPs predict SLE in this cohort through a combination of xenobiotic-metabolism variants, a few nominal immune and folate effects and age, with a substantial share coming from SNPs of uncertain genotype quality.

### What the leakage analysis shows

Fitting ComBat inside each fold and keeping each individual in a single fold barely changed the cross-validated scores, and with shuffled labels the full pipeline scored at chance. For this data set, procedural leakage was not the main source of optimism in the earlier analysis. This should not be read as evidence that the pooled numbers are reliable. The batch-only baseline shows that when disease labels are unevenly spread across studies, a classifier can reach a high pooled AUC by learning study identity, even when every preprocessing step is fitted on training data only. Mean-only ComBat with a disease covariate removes differences in batch location while protecting disease differences, but it does not remove differences in variance, covariance or tissue composition between, for example, a whole-blood cohort and a sorted-cell cohort, and a flexible model such as XGBoost can still pick these up. The within-batch and leave-one-batch-out results are therefore the more realistic estimates of how well these 11 genes generalise.

### Why PR-AUC and MCC matter here

The expression data are 70.6% SLE, and one batch is 93.6% SLE. In that setting a classifier that calls nearly every sample SLE still obtains a PRAUC close to 1 and a reasonable accuracy and F1. MCC was the only metric that exposed this behaviour (MCC 0.009 for L1-LR in GSE45291), and it also separated the models more clearly than ROC-AUC did (Table 6). For imbalanced biomedical classifiers, PR-AUC should be reported together with its chance level, alongside MCC.

### Limitations

The genotype cohort is ethnically restricted (South Indian), limiting generalisability, and no independent genotype cohort was available for replication. Four SNPs deviate from HWE in controls and the strongest association shows a pattern typical of genotyping error; the alleles behind the genotype codes were not available, so effect directions cannot be tied to specific alleles; and controls were on average four years older than cases, so age contributes much of the SNP models’ discrimination. The candidate-gene design of both data sets introduces ascertainment bias, and variants outside these pathways are invisible. Only 11 genes were available after filtering, which limits the empirical Bayes shrinkage in ComBat and is why full ComBat was unstable. Disease is partly confounded with batch: two batches have no SLE samples, one has no controls, and one batch contributes most of the SLE samples. In such designs batch adjustment can bias group differences (Nygaard et al., 2016), and some of the remaining between-study heterogeneity reflects real differences in cohort, tissue or platform rather than technical batch. The two centred series could not be placed on the log_2_ scale, so cross-platform comparability relies on the location adjustment in ComBat. Subject identities were inferred from the GEO sample titles, and samples whose title did not identify the person were assumed to come from different individuals. Some healthy-control samples in GSE38351 and GSE39088 were stimulated *in vitro* (for example with interferons) and may not represent resting healthy blood. Finally, the lupus nephritis biopsies in GSE32591 were kept under their curated Other/Unknown label and were not used as SLE cases.

## Conclusion

Pathway-level candidate-gene genotyping yields modest SLE discrimination (ROC-AUC *≈* 0.70, MCC *≈* 0.31), and a substantial part of even this modest signal depends on SNPs that fail HWE in controls. The most credible genetic signals point to xenobiotic metabolism (*CYP1A1* M2 and M4, *COMT* and *GST T1*). The same pathway genes measured at the transcriptomic level discriminate better, but when samples are pooled across public studies, much of the apparent advantage comes from study membership rather than disease biology. Leakage-free cross-validation with fold-wise batch correction should be standard for such data, but it is not enough on its own: batch-only baselines, within-study metrics, held-out-study validation and permutation checks should be reported alongside pooled scores. Reduced *MTR* expression is the most robust molecular finding, consistent across model importance, within-study comparisons and batch-adjusted testing, and is a candidate for validation in a cohort where genotype and expression are measured in the same individuals.

## Data and code availability

The curated data, all analysis code and every table and figure in this paper are available at https://github.com/catdisk04/AIBD_project. Running run_all.sh regenerates all results from the input data in about five minutes. The scripts in the code/ folder produce the following:

- 01_batch_correction.py: Tables 1 and 2, Fig. 1;
- 02_gex_classification.py: Tables 6, 7, 8 and S1, Fig. 3;
- 03_snp_analysis.py: Tables 5, 3 and 4, Fig. 2;
- 04_differential_expression.py: Table S2, Fig. 5;
- 05_figures.py: Figs. 4 and 6;
- 06_sanity_checks.py: the permutation checks and Fig. S1.

Analyses used Python 3.13 with NumPy, pandas, SciPy, statsmodels 0.15, scikit-learn 1.9, XGBoost 3.4 and SHAP 0.54, with fixed random seeds throughout.

## Author contributions

- **Vasudha Kannan:** Carried out the preprocessing, normalisation and batch correction of the gene-expression data and provided the final cleaned data set for analysis.
- **R Karthik:** Team leader (coordinated task allocation, scheduled meetings and set the overall plan of action); identified the relevant data sets, contributed to the later stages of the analysis and prepared the manuscript.
- **Aldis Daniel:** Conducted the literature survey, helped in choosing the problem statement and took part in the manual curation of the gene-expression data set.
- **Hitesh Kandarpa:** Designed and implemented the analysis pipeline.

All authors contributed equally in terms of time spent on the project and approved the final manuscript.

## Acknowledgements

We thank Prof. Meiyappan Lakshmanan, Indian Institute of Technology Madras, for his guidance and for his feedback on this work.

## Supplementary material

**Figure S1:**
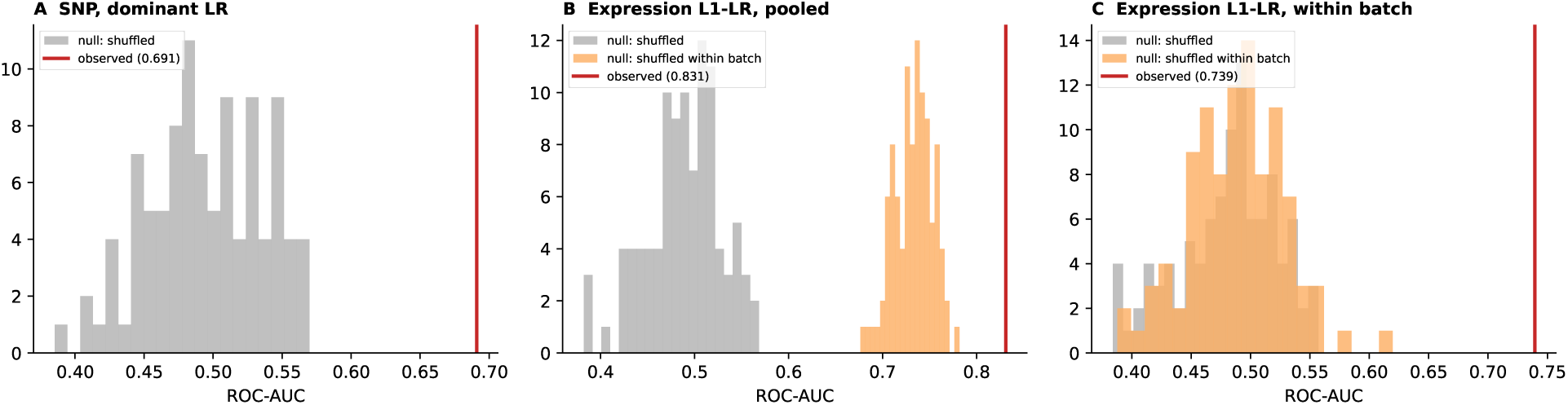
Label-permutation checks (100 permutations each, first cross-validation split). (A) SNP dominant LR with labels shuffled across subjects. (B, C) Leakage-free expression L1-LR, pooled and within-batch ROC-AUC, with labels shuffled across all samples (grey) or only within batches (orange). Red lines mark the observed values.

**Table S1:** Leave-one-batch-out validation. The held-out batch is shifted onto the training-set gene means without using its labels.

| Held-out batch | *n* | SLE | L1-LR |  |  | XGBoost |  |  |
| --- | --- | --- | --- | --- | --- | --- | --- | --- |
|  |  |  | ROC | PR | MCC | ROC | PR | MCC |
| GSE10325_GPL96 | 67 | 39 | 0.553 | 0.629 | 0.071 | 0.463 | 0.529 | 0.028 |
| GSE38351_GPL96 | 26 | 14 | 0.554 | 0.658 | -0.181 | 0.482 | 0.531 | -0.098 |
| GSE39088_GPL570 | 142 | 78 | 0.596 | 0.629 | 0.057 | 0.531 | 0.608 | -0.087 |
| GSE45291_GPL13158 | 312 | 292 | 0.684 | 0.973 | 0.141 | 0.741 | 0.977 | 0.153 |

**Table S2:** Batch-adjusted differential expression, SLE vs. control (OLS, expression *∼* SLE + batch, subject-clustered standard errors).

| Gene | $\log_2\text{FC}$ | 95% CI | $p$ | $q$ (BH) |
| --- | --- | --- | --- | --- |
| *MTR* | -0.315 | $[-0.451, -0.180]$ | $5 \times 10^{-6}$ | $6 \times 10^{-5}$ |
| *TYMS* | +0.339 | $[0.177, 0.502]$ | $4 \times 10^{-5}$ | $2 \times 10^{-4}$ |
| *STAT4* | -0.238 | $[-0.378, -0.097]$ | 0.0009 | 0.0033 |
| *GSTP1* | -0.183 | $[-0.308, -0.058]$ | 0.0042 | 0.0116 |
| *TLR2* | +0.178 | $[0.019, 0.338]$ | 0.0283 | 0.0623 |
| *GSTT1* | -0.107 | $[-0.315, 0.102]$ | 0.317 | 0.513 |
| *TLR4* | +0.097 | $[-0.105, 0.300]$ | 0.347 | 0.513 |
| *MTRR* | -0.040 | $[-0.139, 0.059]$ | 0.424 | 0.513 |
| *TLR7* | +0.089 | $[-0.142, 0.319]$ | 0.452 | 0.513 |
| *CYP1A1* | +0.042 | $[-0.071, 0.154]$ | 0.466 | 0.513 |
| *MTHFR* | +0.032 | $[-0.063, 0.127]$ | 0.514 | 0.514 |

