## Supplementary figures and images for "Prediction of Systemic Lupus Erythematosus from Genotype and Gene-Expression Data: Batch Correction and Leakage-Free Evaluation"

### Supplemental figure 1

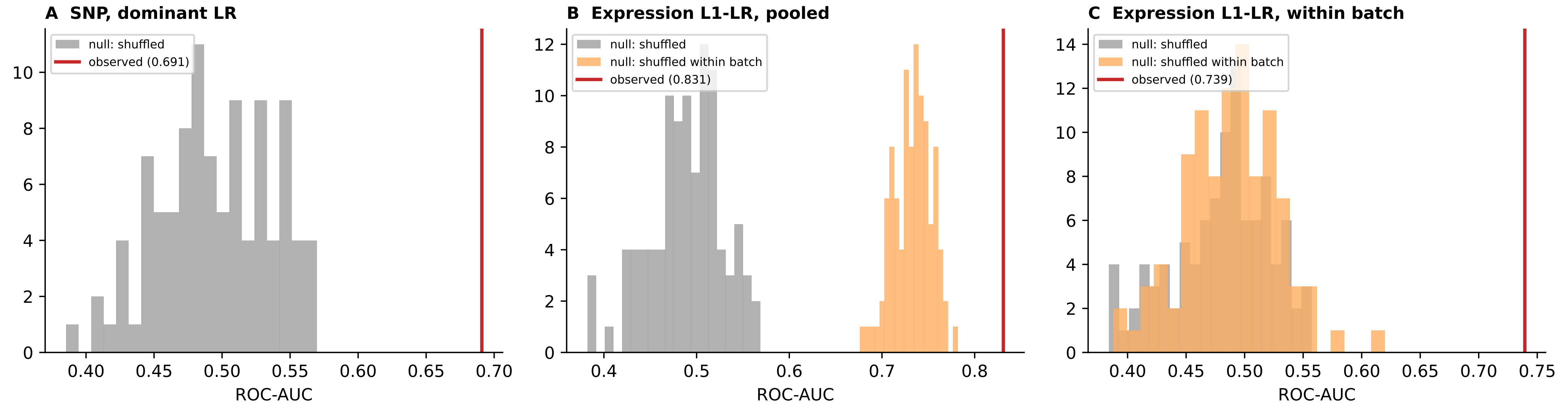
